# High-salt diet disturbs the physiological uterine endothelial junction remodeling during pregnancy

**DOI:** 10.1101/2025.11.06.687038

**Authors:** Lin Huang, Shuangbo Kong, Qionghua Chen, Ziying Huang, Min Li, Ning Guo, Xinran Yu, Minshan Huang, Haibin Wang, Dunjin Chen, Jingsi Chen

## Abstract

A high-salt diet (HSD) induced excessive sodium intake is a major risk factor for various diseases. However, its specific effects on uterine endothelial cell and pregnancy outcomes remain poorly understood. In this study, we demonstrated that HSD induces gestation specific hypertension and adverse pregnancy outcomes in murine models. The proper spiral artery remodeling, which involved the decidual vascular endothelia, is thought to be essential for normal regulation of gestational blood pressure. We explored the molecular basis for the adaptation of vascular endothelial cell based on the transcriptome data, and uncovered the significant changes for the cell junction in the endothelia during this process. Furthermore, it was revealed that HSD induced more salt accumulation in the uteri and triggered the disturbed vascular endothelium cytoskeletal remodeling, which was associated impaired trophoblast cell invasion and incorporation into the vascular wall. Overall, our study identifies unique roles of decidual endothelium for pregnancy vascular remodeling, and demonstrates that HSD disrupts physiological spiral artery remodeling through endothelial maladaptation, which may concurrently induce systemic endothelial damage in maternal systemic vascular.

## Introduction

The World Health Organization (WHO) recommends salt intake of less than 5 g per day for adults and even less for children (1). As socioeconomic development progresses, human dietary habits have changed resulting in increased food intake, increased use of food additives and diets high in fat and sugar. Consequently, daily salt consumption is far higher than WHO recommendation particularly in developed and developing countries. Nowadays strong evidence proves that excess salt intake contributes to many diseases and significantly increases cardiovascular morbidity and mortality (2, 3). Low-salt diets can reduce blood pressure in middle-aged and older individuals, irrespective of antihypertensive medication, race or baseline blood pressure (4). High salt-diet (HSD) plays a significant role in many diseases, such as accelerating chronic kidney disease progression, promoting the progression of rheumatoid arthritis, asthma, psoriasis, weakening the immune system and increasing the risk of type 2 diabetes (5–9). Maternal taste changes during pregnancy, including increased cravings for salt (10). However, there are no specific salt intake recommendations for pregnant women, and few studies have reported the association between excessive salt intake and adverse pregnancy outcomes (11).

In this study, we established a high-salt intake mouse model to mimic the human daily HSD state and investigate its effects on pregnancy outcomes. It was revealed that HSD induced unexpected and serious adverse pregnancy outcomes, accompanied with gestation specific hypertension. The proper spiral artery (SA) remodeling, accompanied with replacement of decidual endothelial cell (EC) by invasive trophoblast, is thought to be essential for normal regulation of gestational blood pressure. We performed transcriptomic profiling to explore the decidual EC adaptation during the pregnancy progression, and uncovered the significant endothelial cytoskeletal remodeling with loosen endothelial cell-cell junctions. Furthermore, we found that HSD resulted in sodium accumulation in the uterine decidua, and demonstrated that high salt microenvironment in the decidua caused disturbed endothelial adaptation, which interrupts SA remodeling process and adversely impacts pregnancy outcome.

## Results

### HSD enhances maternal BP and causes adverse pregnancy outcome

We first established a mouse model fed a high-salt diet (HSD, 2% NaCl in drinking water, Figure 1A). Compared to the control group (CON), HSD female mice exhibited a significant increase in water consumption, while food intake and body growth remained unaffected after long-term HSD exposure in non-pregnant females (Supplemental Figure 1, A-C). Continuous eleven-week monitoring of both non-pregnant and pregnant mice (day 18 of gestation, D18) revealed no significant effect of HSD on body weight or apparent water retention, as demonstrated by MRI-based body composition analysis (Supplemental Figure 1F). Urinary protein-to-creatinine ratio measurements and histological assessment via Masson staining indicated no overt structural or functional renal impairment in HSD mice (Supplemental Figure 1, H, I and J). Excessive salt intake is one known important risk factor for many diseases, with most cardiovascular diseases associated with HSD attributable to hypertension (3, 12–15).

**Figure 1.**
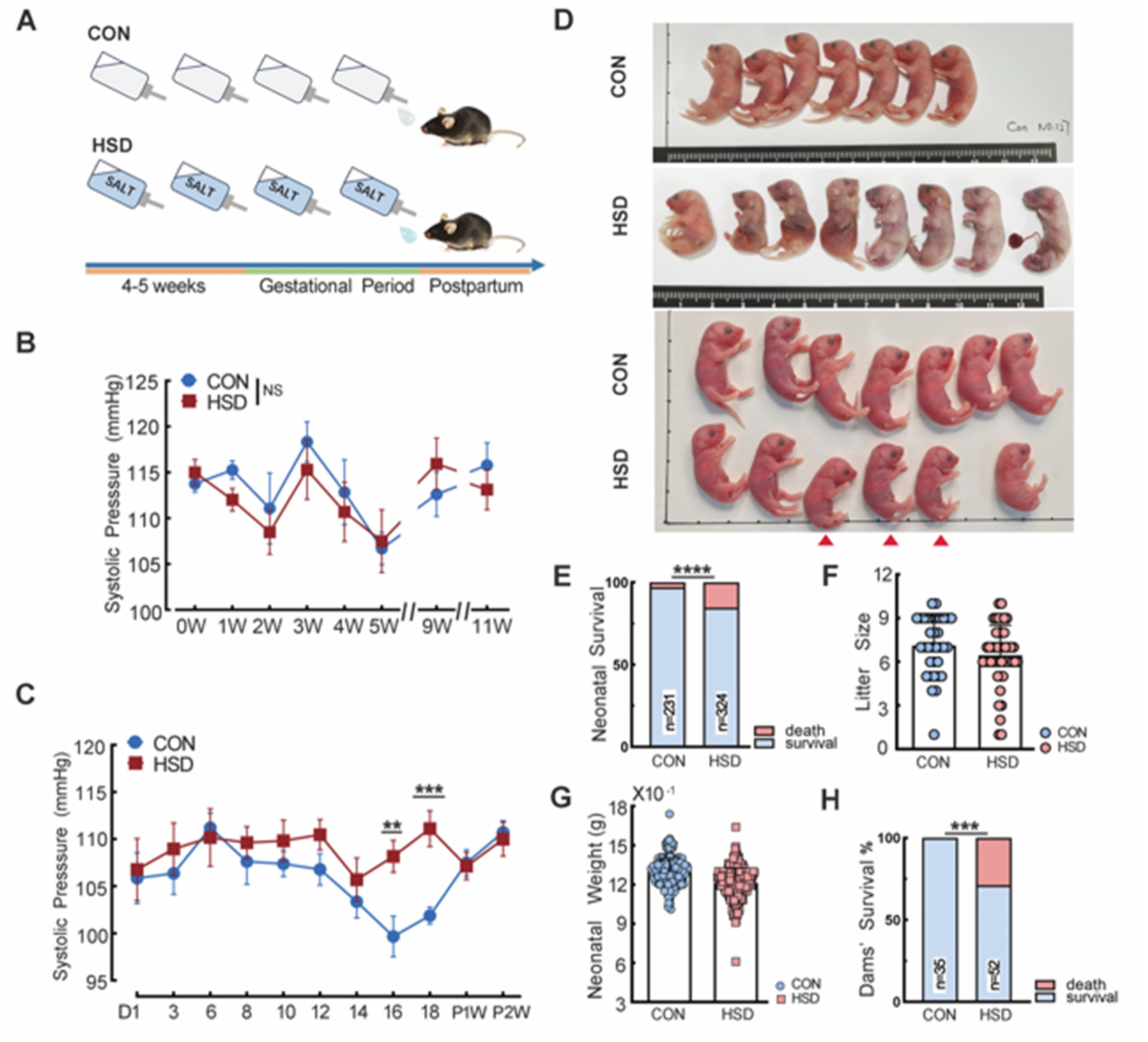
HSD enhances maternal systolic blood pressure (BP) and causes adverse pregnancy outcome. (A) Schematic diagram representing experimental approach of normal salt diet (CON) and high-salt diet (HSD) mice model. Aged 6-7 weeks old female mice feeding with respective drinking water for one month before mating until postpartum one month. (B) HSD did not impact non-pregnant female’s mice systolic BP. Unpaired *t* tests. data are presented as Means ± SEM (n = 5-to-10 animals in CON; n = 6-to-11 animals in HSD). NS, no significant. (C) Blood pressure measured by noninvasive tail-cuff system (n = 9-to-15 animals per group). Unpaired *t* tests. data are presented as Mean ± SEM. D16: \*\**P* = 0.005, D18: \*\*\**P* = 0.0003. (D) Representative image of pregnant outcome. The pictures show the normal phenotype of CON compared to HSD neonatal mice, which exhibit (top) stillbirth and (bottom) smaller postnatal mice (red arrows). (E) Neonatal mice survival in postnatal first day. 224 over 231 animals in CON; 275 over 324 animals in HSD. Chi-square tests. \*\*\*\**P* < 0.0001. (F) Number of pups per litter. n = 34 dams in CON; n = 46 dams in HSD. Data are presented as Mean ± SD. Unpaired *t* tests. NS = no significant. (G) Offspring body weight at postnatal day one. n = 224 neonatal mice in CON; n = 274 neonatal mice in HSD. Data are presented as Mean ± SD. Unpaired *t* tests. \*\*\*\**P* < 0.0001. (H) Percentage of dead dams died within postpartum one month. (0 over 35 in CON; 15 over 52 in HSD). Fisher’s exact test. \*\*\**P =* 0.0002.

Therefore, we investigated the impact of HSD on maternal blood pressure (BP) by noninvasive tail-cuff system. Weekly BP measurements in non-pregnant females confirmed that high-salt diet in our study causes systolic blood pressure differences between groups, and this was exclusively observed in pregnant mice (Figure 1B). While once pregnant, BP rose from the first day of gestation day (day 1, D1) to a peak at D6, followed by a gradual decline until D16 in CON group mice (Figure 1C). This progressive BP reduction during gestation aligns with previous reports (16–20). In contrast, HSD mice displayed sustained BP elevation from D1 to D12, with a transient decrease between D12 and D14, followed by a renewed increase to peak levels at D18 (peak, *P*<0.05). Notably, HSD dams delivered smaller offspring with elevated stillbirth rates upon observation on the first postpartum day (Figure 1D, 1E and Supplemental Figure 2A). Pathological examination revealed that many stillborn fetuses from HSD dams exhibited full developmental maturity, suggesting late-gestation demise. Neonates from HSD dams demonstrated significantly reduced body size and weight at birth but without effect on litter size, with an average 6.56% weight reduction compared to CON neonates (Figure 1, F and G), suggesting an intrauterine growth restriction (IUGR) phenotype. Strikingly, HSD dams exhibited significantly higher maternal mortality within one month postpartum (Figure 1H). Collectively, these severely adverse outcome of pregnancy-specific hypertension following the HSD treatment suggested that the SA remodeling may present defects (21, 22).

### HSD leads to adverse pregnant outcome and defective SA remodeling

Stillborn fetuses carried by HSD dams exhibited developmental abnormalities and placental dysplasia (Supplemental Figure 2, A and B). HSD group displayed a significantly sparser placental vasculature compared to the vascular network of CON, based on the Three-dimensional (3D) visualization of D18 placentas (stained with CD31 antibody), suggesting impaired placental development in HSD (Figure 2A). Spiral artery remodeling, a critical process during placental development, involves trophoblast cell invasion into the inner layer of SA vessels, displacing endothelial cells and smooth muscle to enhance uterine perfusion and placental blood flow (23, 24). Therefore, we investigated whether HSD disrupted SA remodeling.

**Figure 2.**
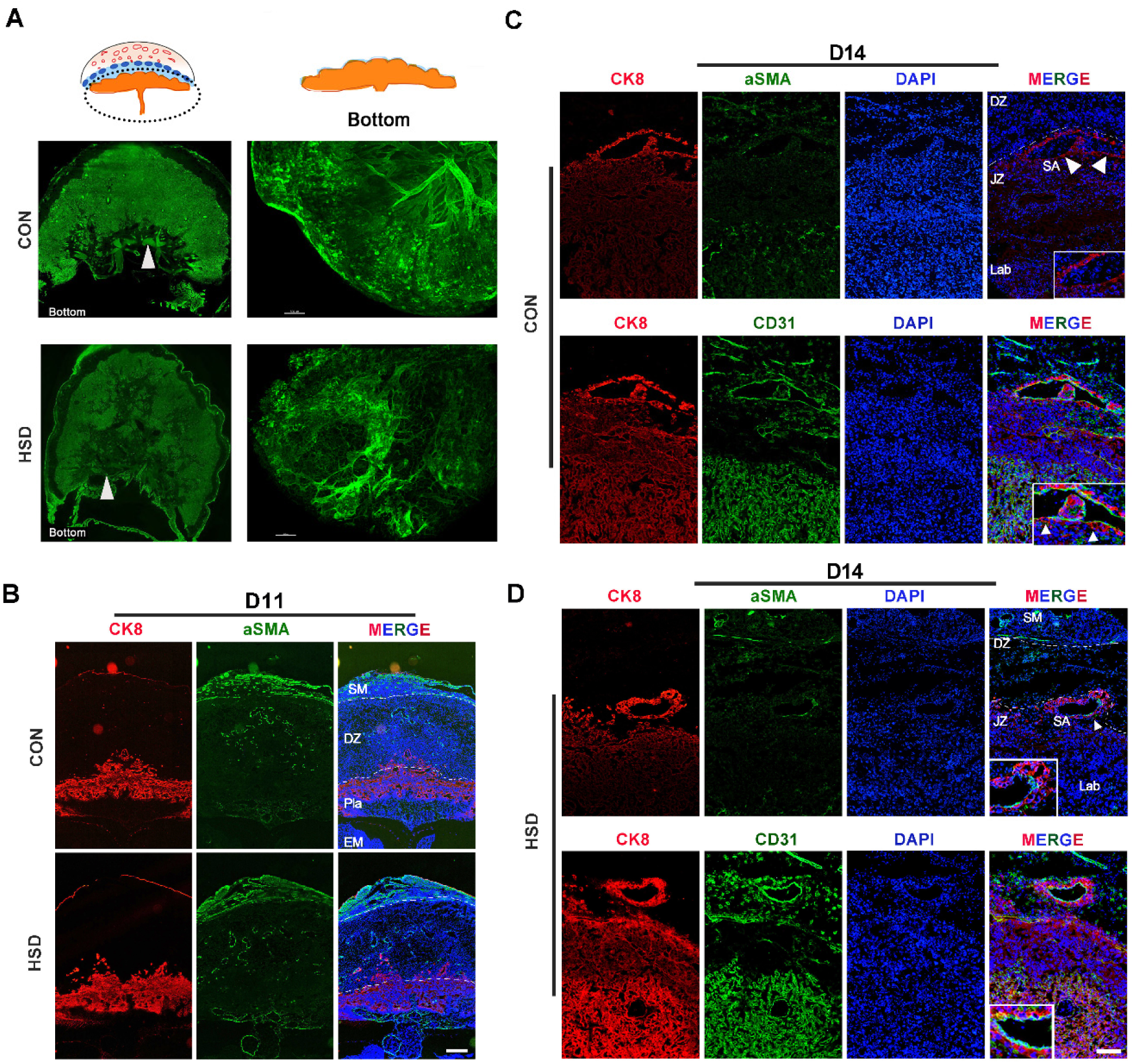
HSD lead to defects in placentation and spiral artery remodeling. (A) 3D visualization of D18 placenta vascular tree. As shown in the representative photos, labyrinthine vascular net (at the bottom of placenta) of HSD is sparser than CON. scale bars, 500 μm. (B) Trophoblast cells invade into decidua and smooth muscle still presents in spiral arteries in D11. SM, smooth muscle layer. DZ, decidual zone. Lab, labyrinth. EM, embryo. scale bars, 500 μm. (C and D) HSD defects in spiral artery remodeling were detected using immunohistochemistry by D14 frozen tissues: Invading trophoblasts displaced ECs in CON. HSD retain tight ECs connection (green), more SMs of SA coverage (green) and TBs are totally blocked outside ECs. Longitudinal sections: (top) were stained for CK8 (cytokeratin-8) (red) for trophoblasts and αSMA (green) for smooth muscle cells, (bottom) were stained for cytokeratin (CK 8) (red) for trophoblasts and CD31 (green) for endothelial cells. Nuclei were stained with DAPI. SA, spiral arteries. JZ, junctional zone. Arrows, SA (spiral arteries). scale bars = 100 μm. Every experiment has more than 3 biological replications.

At D11, initial trophoblast cell invasion into the decidua showed comparable progression between two groups (Figure 2B). By D14 morning in CON placentas, serial section of placenta showed that trophoblast cells were surrounding the SAs and the middle smooth muscle layer was near-complete loss (Figure 2C, top), with disorganized endothelial cells in the inner layer and multiple trophoblast cells infiltrating the SA lumen (Figure 2C, bottom). In contrast, HSD placentas retained a thicker smooth muscle layer despite trophoblast cell encirclement of SAs (Figure 2D, top). Strikingly, the endothelial layer maintained structural integrity in HSD mice, effectively blocking further penetration of trophoblast cell (Figure 2D, bottom). These observations were corroborated by colocalization analysis of endothelial, smooth muscle, and trophoblast cell markers (Supplemental Figure 2D). This raises the question of whether SA’s endothelial adaptations are required to permit trophoblast invasion during SA remodeling.

### ECs of SA display adaptations during pregnancy progression

To address this question, we performed bulk RNA-sequencing (RNA-seq) transcriptional profiling of vascular endothelial cells in decidua during the pregnancy progression. Firstly, maternal decidua tissues, isolated from placental via mechanical dissection under microscopic guidance, were digested with collagenase to get the single cell suspension, and endothelial cells were purified by flow cytometry through CD31 staining prior to bulk RNA-seq (Figure 3A). Analysis of endothelial RNA-seq data notably revealed dynamic activation of decidual endothelium during SA remodeling. Principal component analysis (PCA) demonstrated distinct transcriptional profiles across placental development stages (D8, D11, D14), with D14 samples forming the most distinct cluster (Figure 3B). Transcriptional divergence was most pronounced between D11 and D14. Differential gene expression analysis and hierarchical clustering of CON endothelial transcriptomes identified nine core expression patterns spanning D8 to D14 (Figure 3C). Notably, multiple clusters exhibited dramatic expression shifts between D11 and D14 (e.g., contrasting trends in clusters 3 vs 7 and 9; 1 vs 6) (Figure 3C). Based on these findings, we focused on physiological characterizing of endothelial adaptations during the D11-to-D14 transition— also it is the critical window for spiral artery remodeling.

**Figure 3.**
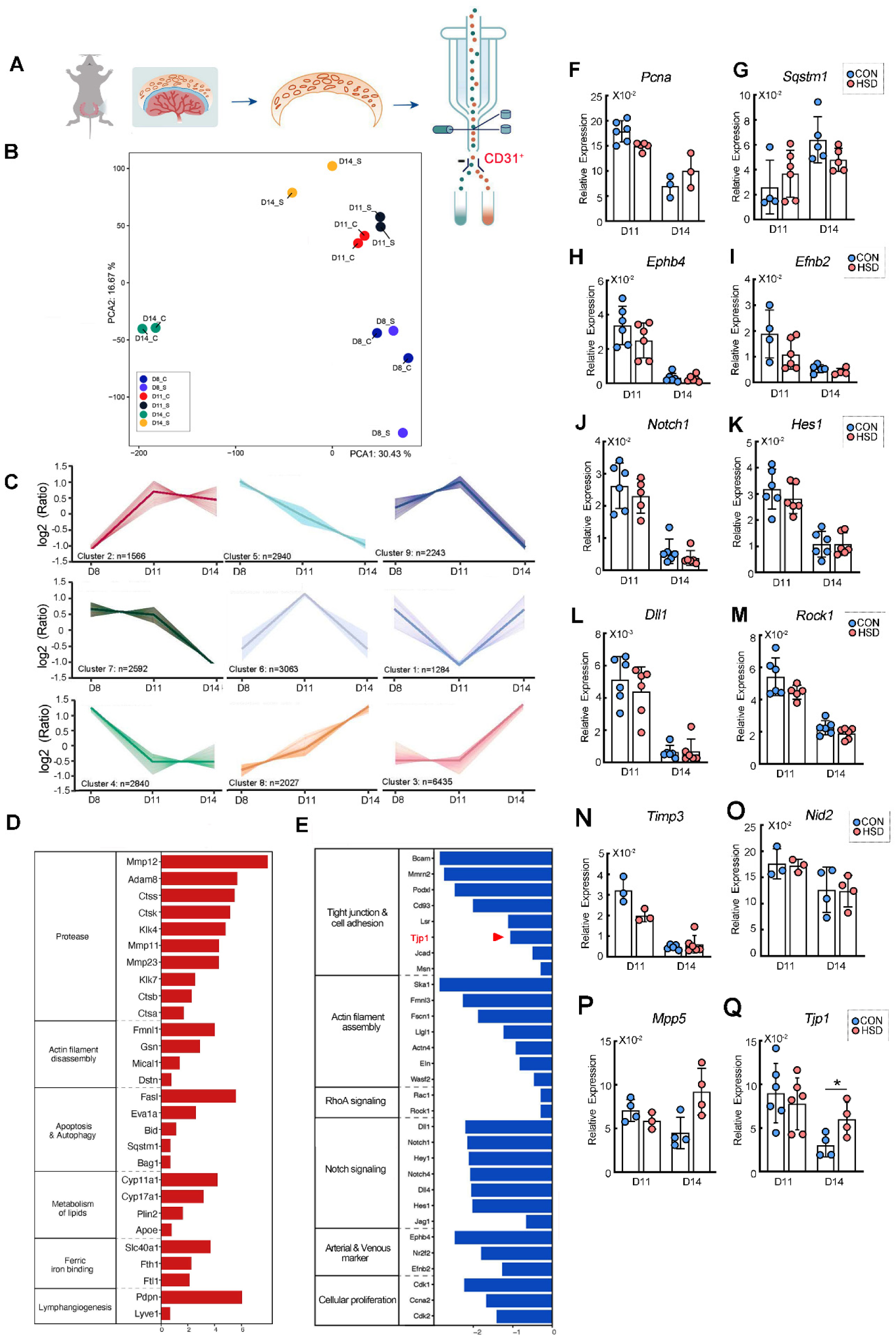
Decidua endothelial cells undergo significant gene expression change during remodeling. (A) Schematic depicting study design for RNA sequencing. ECs was isolated by physical separation decidual tissue from living individuals, digested into single cell and collected by flow cytometry for last bulk RNA-sequencing transcriptional profiles. (B) PCA based on transcriptomics data of ECs from bulk RNA-Seq of CON and HSD groups reveal the distribution of D8, D11 and D14. (n = 3-to-5 animals per group). D8-C, D11-C and D14-C represent CON’s uterus decidual endothelium cells in D8, D11 and D14 respect. D8-S, D11-S and D14-S represent HSD’s uterus decidual endothelium cells in D8, D11 and D14 respect. (C) Different type of change trend of CON’s genes in bulk RNA-sequencing transcriptional profiles. (D) The upregulated genes enriched in CON from D11-to-D14. (E) The downregulated genes enriched in CON from D11-to-D14. (F-Q) Quantitative RT-PCR analysis of gene expression of CON and HSD. The gene expression was normalized to GADPH. Mean ± SEM. Data are shown as more than 3 biologically independent replicates. \**P* < 0.05.

Next, we focused on D11-to-D14 gene expression analysis. These genes were enriched in a variety of cellular processes, including upregulation of mRNAs encoding molecules associated with cellular apoptosis and autophagy (*Fasl, Sqstm1*), actin filament disassembly (*Fmnl1, Gsn*) and lymph angiogenesis (*Pdpn, Lyve1*) (Figure 3E). Notably, downregulated genes were implicated in cellular junction (e.g., B*cam, Mmrn2, Tjp1*), actin assembly (e.g., *Fmnl3, Fscn1, Wasf2*), arterial and venous marker (e.g., *Ephb4, Efnb2*), and genes involved in Notch and RhoA/ROCK signaling pathways—critical regulators of angiogenesis and cytoskeletal dynamics, respectively (25) (Figure 3D). To validate these findings, we conducted RT-PCR and immunofluorescence assays (Figure 3, F-Q). Intriguingly, HSD mice exhibited marked transcriptional divergence in key genes (e.g., *Mpp5* and *Tjp1*) at D14 (Figures 3, P and Q). *Mpp5*, a tight junction-associated protein (26), showed particularly pronounced differential expression. Co-staining of Ki67 and CD31 revealed significantly reduced endothelial proliferation from D11 to D14 in both groups (Supplemental Figure 3A). Collectively, our RNA-seq data demonstrate that vascular endothelium in decidua undergoes functional reprogramming during SA remodeling, characterized by: (a) Cessation of cellular proliferation; (b) Enhanced apoptotic activity; (c) Lymphangiogenic mimicry; (d) Loss of junctional integrity—potentially mediated through cytoskeletal remodeling and reduced cell-cell adhesion. A critical intergroup difference also emerged that CON SA endothelium exhibited greater structural loosening compared to HSD during remodeling. This observation raises a pivotal question—could defective endothelium adaptational changes in HSD impair trophoblast invasion and displacement. To investigate this, then we compared D14 transcriptional profiles between groups.

### HSD causes defective cytoskeleton remodeling in EC and blocks SA remodeling

RNA-seq transcriptome analysis revealed up-regulation of known cytoskeletal actin assembly, maintain extracellular matrix structure, vascular endothelial cadherin and cellular junction genes such as Eln (elastin), Rasip1 (ras interacting protein 1), Fmnl3 (formin-like 3), Cdh5 (cadherin 5) and Tjp1 (tight junction protein 1) (27–29) in HSD group. Gene Ontology (GO) enrichment analysis of different expressed genes on D14 endothelia revealed significant enrichment in cell-substrate junctions, actin filament regulation, and tight junction pathways (Figure 4B). Subsequent Gene Set Enrichment Analysis (GSEA) of cellular biological states yielded further present our findings (Figure 4C-F). In the “Apical Part of Cell” category, differentially expressed genes were linked to enhanced cellular polarity, adhesion maintenance, and actin cytoskeletal remodeling, including upregulated *Vangl2*, *Fat4*, *Tjp1*, and Notch pathway activation (25, 30–34). In the “Extracellular Matrix Structural Constituent” category, HSD mice exhibited marked upregulation of ECM-related genes *(Ecm1, Fbln5, Lama1/3/4, Lamb2).* We compiled a set of genes encoding tight junction, cell adhesion and actin filament assembly across D11–D14. While both groups showed similar downregulation trends, the CON group exhibited more pronounced downregulation compared to HSD (Figure 4G).

**Figure 4.**
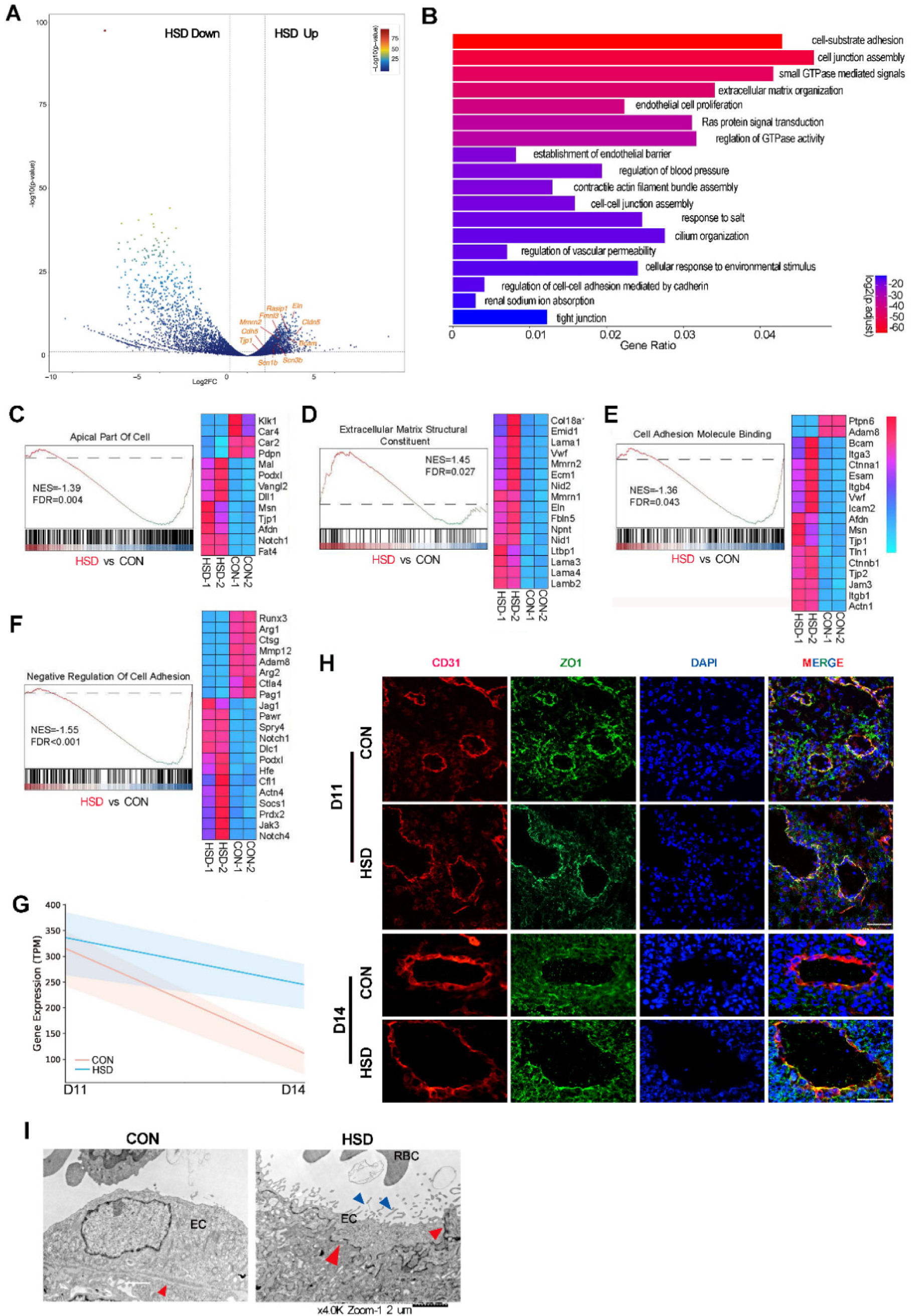
HSD disrupts spiral artery endothelial cells remodeling. (A) Volcano plot showing differentially expressed D14 endothelium genes between CON and HSD. *P* values determined via *T* test. (B) GO enrichment analysis: RNA-sequencing reveals defects in the endothelial junction adaptational change in HSD. (C-F) Gene set enrichment analysis (GSEA) results showing. (G) Overall gene change trend in cellular junction, adhesion and actin filament. (H) Immunofluorescence images showing spiral artery endothelial tight junction become loosen from D11-to-D14 in CON but disrupt in HSD. Scale bars = 100 μm. (I) Transmission electron microscopy images of spiral artery endothelium. scale bars = 2 μm. EC, spiral artery endothelial cells. RBC, maternal red blood cells in spiral artery (red arrows, endothelial junction of cell-cell junction and endothelial cell-extracellular matrix junction. blue arrows, apical microvilli). Every experiment has more than 3 biological replications.

Tight junction was focused as a cellualr event for endothelial adaptation. Although RNA-seq data encompassed all decidual vascular endothelia, we confirmed that SA’s endothelium presents this characteristic by co-immunofluorescent. ZO1 (Tjp1) localized robustly and displayed stronger between SA endothelial cells’ boundary at D11 in both groups. By D14, ZO1 signal was nearly absent in CON, whereas HSD retained detectable expression (Figure 4H). Electron microscopy further corroborated this divergence (Figure 4I). We also observe that HSD endothelia displayed apical microvilli structures, which was critical for cytoskeletal linkage, polarity maintenance and hemodynamic resistance modulation (Figure 4I). (35–39).

### Sodium accumulates in decidua and EC cytoskeletal rearranges in high salt microenvironment

Prior evidence has proved the Na⁺ deposition in skin, muscle, and endothelial glycocalyx under HSD. This prompted us to assess uterine sodium accumulation in HSD during pregnancy. Serum sodium analysis via ion-selective electrode (ISE) revealed physiological dilution of Na⁺ from D11 to D14, with no significant intergroup difference (Figure 5A). However, inductively coupled plasma optical emission spectroscopy (ICP-OES) of D18 decidual tissue demonstrated significantly Na⁺ accumulation in HSD compared to the CON’s (*P* < 0.05, Figure 5B), indicating sustained sodium retention in the decidua with long term high salt intake. To investigate the mechanistic impact of this sodium-enriched microenvironment on endothelial function, we cultured human umbilical vein endothelial cell (HUVEC) in high-salt medium (HS) to mimic in vivo conditions. HS-treated cells exhibited cortical F-actin redistribution—forming a peripheral ring-like structure (Figure 5C). To rule out the possible osmotic effects, we applied iso-osmotic urea (equal to 40mM HS), and uncovered that it failed to induce F-actin meshwork accumulation and increase junctional ZO1 intensification (Figure 5D). Hence, we wondered whether cortical actin meshwork changes response to the intracellular Na+ increasement. We employed the Na⁺-sensitive probe SBFI-AM to assess intracellular sodium flux, revealing markedly elevated cytosolic Na⁺ in HS-treated cells (Figure 5E). This aligns with known Na⁺-dependent actin remodeling via cation-binding motifs (40). We further hypothesized that Na⁺ influx modulates cytoskeletal reorganization and junctional reinforcement. Amiloride (Na⁺ channel inhibitor) was applied and it can abrogate HS-induced cortical F-actin accumulates beneath submembrane even under extreme hypernatremia (Figure 5F). Given the RhoA/ROCK pathway’s central role in cytoskeletal regulation (41). We inhibited ROCK with Y27632. This treatment effectively blocked HS-driven actin remodeling either.

**Figure 5.**
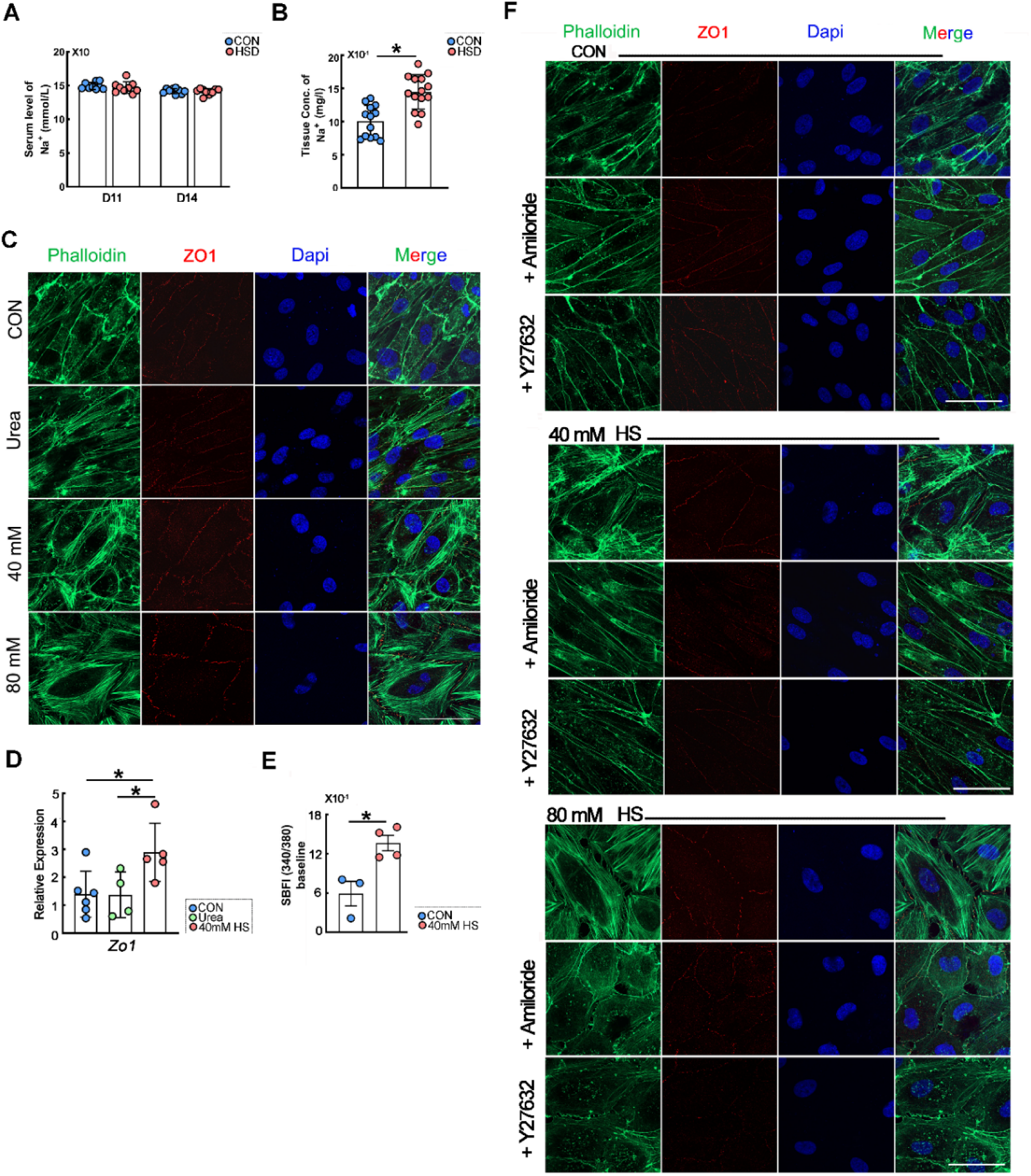
HSD accumulates salt in decidua and EC cytoskeletal rearranges in high salt microenvironment. (A) Serum Na^+^ concentration of gestational D11 and D14. (B) Decidual tissue Na^+^ content determined by inductively coupled plasma-optical emission spectrometry (ICP-OES). Mean±SEM. Data are shown as more than 3 biologically independent replicates. \**P* < 0.05. (C) High salt cell culture medium (HS) impact endothelial cellular cytoskeletal but not case by osmotic pressure. (D) HUVECs quantitative RT-PCR analysis of *Zo1* expression. The gene expression was normalized to GADPH. (E) HUVECs intracellular Na^+^ content by indicator SBFI-AM. One dot presents 1/96-well plate (4000-5000 per well). (F) Amiloride and Y27632 inhibit HS-induced F-actin accumulation. Data are shown as more than 3 independent replicates. scale bars = 50 μm

## Discussion

High-salt diets (HSD) have long been implicated in cardiovascular pathologies, diabetes progression, renal infection exacerbation, and hypertrophic scarring, though recent studies also suggest paradoxical benefits in tumor suppression and vascular barrier enhancement (9, 42–44). However, the association between excessive salt intake and adverse pregnancy outcomes remains underexplored. Our study provides the evidence that HSD induced impaired remodeling of spiral artery (SA). We demonstrate that HSD triggers decidual sodium accumulation, mechanistically impeding SA endothelial remodeling of tight junction integrity, which occurred for vascular adaptation during the normal pregnancy. Notably, our HSD mouse model recapitulates subclinical human dietary salt overconsumption, exhibiting no overt systemic perturbations in non-pregnant state. Consistent with physiological hemodynamic adaptations in pregnancy—increased cardiac output, expanded plasma volume, and reduced peripheral resistance culminating in mid-gestational blood pressure nadir (45). CON mice mirrored established blood pressure trends (Figure 1B). In contrast, HSD dams displayed sustained systolic hypertension (Figure 1B), fetal developmental anomalies, and elevated maternal postpartum mortality (Figure 1). Placental 3D imaging revealed hypovascularization of vascular network and in HSD (Figure 2A), coupled with defective SA remodeling (Figures 2C and D). From endothelial cell aspects for SA remodeling, we uncovered many adaptational changes to support junctional loosening for facilitating trophoblast invasion. HSD disrupted this plasticity, preserving endothelial tight junctions, apical microvilli, and polarity (Figure 4). We propose that HSD reprograms endothelial cytoskeletal dynamics via Na⁺-mediated RhoA/ROCK activation (Figures 5F), diverging from hypertonicity-driven actin assembly (46). Both Na⁺ influx blocked by Amiloride and ROCK inhibitor treatment rescued HS-induced cytoskeletal remodeling phenotypes, suggesting that HSD reprograms endothelial cytoskeletal dynamics via Na⁺-mediated RhoA/ROCK activation (Figures 5F). Notably, decidual Na⁺ retention in decidua under chronic HSD (Figure 5B) extends prior observations of extrarenal sodium storage (17, 42, 47–49). This sodium-enriched microenvironment likely alters endothelial biomechanics as cortical F-actin reinforcement (Figure 5C) enhances junctional tension(50), impeding trophoblast-driven endothelial displacement.

Preeclampsia (PE) pathogenesis centers on SA remodeling failure and placental hypoperfusion (51). While prior research mainly focused on trophoblast cells or immunity cells in uterus, our work highlights endothelial maladaptation as a novel contributor. One important part of our work is to reveal important physiological adaptation of decidual vessel endothelium present in spiral artery remodeling process which are not realized and underestimate but play vital role in spiral remodeling process. For example, we have found MMPs significant increasing from D11 to D14. MMPs family has important function in degradation of the extracellular matrix (52). Cytoskeleton “stabled” cells also by bridging cellular inner element to extracellular matrix, such as integrin (53). Integrin family forms the major family of cell adhesion molecules, as a transmembrane protein link between intracellular proteins (indirectly link to the cytoskeleton) and extracellular matrix components (such as fibronectin, laminins and collagens) (54, 55). Defects in extracellular matrix remodeling can cause trophoblast invasion defect (21).

ZO1, one of the primary components of the tight junction locate near the apex of endothelium, binds directly to occluding and connects to the apical and basolateral cytoskeleton in cytoplasm (56–58). The tight junction’s primary functions are to maintain intercellular barrier integrity and preserve cellular polarity (59). In our data, we have found tight junction still kept in high level in D14 of HSD. These observations align with prior reports of vascular barrier stabilization under hyperosmotic NaCl exposure (44). ZO1 can direct interactions with F-actin, and it is the important protein linking the cellular inner actin to outer membrane (60, 61). Alterations to the actin cytoskeleton critically regulate tight junction stability (62). Our findings provide a mechanistic framework for spiral artery endothelial adaptation: junctional loosening facilitates trophoblast invasion and endothelial replacement during SA remodeling. Combing our cellular experiment, this essential plasticity is defective in HSD where cytoskeletal rearrangements reinforce tight junctions under sodium-enriched microenvironmental conditions. Our work establishes endothelial cytoskeletal remodeling as a pivotal biological event in SA remodeling. Increased decidual endothelial rigidity may impair vasodilatory factor release and elevate vascular resistance, contributing to reduced uteroplacental perfusion and IUGR (63). Interestingly, previous studies have reported that exposure of the endothelium to mild hyperosmolarity enhances vascular barrier function (44).

The Interstitum and endothelial glycocalyx network can help buffer and accumulate high amounts of excess Na+, which subsequently inflex into the endothelial cells through Na+ channels and may cause endothelial dysfunction (39). Syndecan-1 (Sdc-1) is one of important member of cellular proteoglycan (64). Inflammatory cytokines, MMPs and oxidative stress can induce syndecan shedding from cellar surface (65, 66). Many diseases have been reported increasing Sdc-1 expression, such as chronic cholestasis, ovarian cancer, invasive breast carcinomas, sterile injuries (67–69). We observed that serum Sdc-1 in HSD mice was significantly higher than CON in D14 and this difference appear earlier than endothelin 1 (ET-1, Supplemental Figure 4, A and B). Elevated serum Sdc-1 mirrors vascular endothelial injury in pregnant hypertension diseases (70, 71), positioning Sdc-1 as a potential biomarker for maternal systemic endothelial dysfunction.

Our findings advocate for prenatal dietary salt guidelines and endothelial-targeted therapies to mitigate SA remodeling defects and protect maternal systemic vascular endothelium during pregnant, especially for those who suffer from gestational hypertension diseases, which is meaningful for the later life of postpartum women.

## Methods

### Mice

Female C57BL/6J mice (6-7 weeks old) were obtained from the Laboratory Animal Center of Xiamen University. The mice were randomly divided into two groups: High-salt diet group (HSD), received 2% NaCl (AR, 10019318) in drinking water; Control group (CON): provided with deionized water. Both groups were maintained on their respective regimens for one month prior to mating with 10-week-old male C57BL/6J mice fed standard chow. All animals were housed under SPF condition with a 12-hour light/dark cycle (08:00-20:00 light phase). Mating pairs were established daily between 16:00-18:00, with vaginal plug checks performed at 08:00-09:00 the following morning (day 1 of gestation, D1). Systolic blood pressure was measured non-invasively using tail-cuff system (Softron Biotechnology, BP-2010A) at 08:00-10:00 daily. All mice underwent three consecutive days of device acclimation prior to formal measurements. This study was conducted in accordance with the NIH Guide for the Care and Use of Laboratory Animals and approved by the Institutional Animal Care and Use Committee of Xiamen University (XMULAC20210105).

### EchoMRI

Body composition analysis (fat and water content) was performed using nuclear magnetic resonance analyzer (EchoMRI™-100H). Each animal underwent triplicate measurements under isoflurane anesthesia, with the average value used for final analysis. All data are reported with more than 3 independent mice.

### Urinary protein and creatinine

Urine samples were collected from non-pregnant females after 8-week dietary intervention, D16-D18 pregnant mice and postpartum day 30 (P30). Samples were immediately stored at -80°C until analysis. Urinary albumin-to-creatinine ratio (ACR) was determined by (creatinine) Sarcosine oxidase method and (albumin) immunoturbidimetric assay. Measurements were performed by biochemical analyzer (Mindray, China). All data are reported with more than 3 independent mice.

### Masson staining

Renal specimens were dissected by removing the surface fibrous capsule, bisected along the coronal plane, and fixed in 4% paraformaldehyde (PFA, Sigma 30525-89-4) for 24 hours at room temperature before tissue processing and embedding. Paraffin-embedded tissue sections (4 μm) were de-paraffinized, rehydrated, and stained with Masson trichrome staining which was performed using a Masson trichrome staining kit and according to the manufacturer’s instructions (Beijing Solarbio, G1346-8).

### Immunofluorescence

Paraffin sections (4μm) were utilized for immunofluorescence (IF) staining, incubated with antibodies: CD31(1:300, R&D), CK8 (1:800, DSHB), α-SMA (1:300, CST), Ki67 (Servicebio, 1:200). Specific secondary antibodies were used to detect the antigen (1:400, Jackson ImmunoResearch), and 4,6-diamidino-2-phenylindole (DAPI) was used to detect the cellular nuclear (1:500, Solarbio).

### RT-qPCR analysis

Total RNA was isolated from cells with TRIzol (Invitrogen) and converted to cDNA (Takara). The cDNA was amplified by RT-qPCR, determine statistical significance by *P* < 0.05. All primers are listed in supplemental material (Table S1).

### Immunostaining for 3D imaging

Placental samples from gestational day 18 (D18) were fixed in Dent’s fixative (methanol: DMSO, 4:1) at -20°C overnight. On the following day, specimens were washed three times with 100% methanol (1 h per wash). To remove pigmentation, tissues were bleached with 3% H₂O₂ (prepared in methanol) at 4°C overnight. After six sequential PBST washes (1 h each), samples were blocked with 5% BSA at 4°C overnight. Primary antibody incubation was performed using anti-CD31 antibody (1:300, R&D) at 4°C for 7 days, followed by species-matched secondary antibody incubation (1:400, Jackson ImmunoResearch) for an additional 7 days. Following dehydration in 100% methanol, tissue clearing was achieved using BABB solution (benzyl alcohol:benzyl benzoate = 1:2) for 30 mins in light-protected condition.

### Flow cytometry

Decidual tissues were mechanically dissociated under microscope and enzymatically digested with collagenase I/V mixture (Worthington, LS004196/LS004188) at 37°C for 60 mins. The enzymatic reaction was quenched with 5% BSA in DPBS. Single-cell suspensions were stained with fluorochrome-conjugated anti-CD31 (1:200; eBioscience, 11031185) and anti-CD45 (1:200; BD Pharmingen, 553081) antibodies at 4°C for 30 min. Flow cytometry analysis was conducted using a Beckman MoFlo Astrios EQS cell sorter. Sorted CD31⁺ cells were immediately lysed in TRIzol reagent (Invitrogen, 15596018) for subsequent RNA sequencing.

### Transmission electron microscopy

Decidual tissues from D18 were dissected to remove the smooth muscle layer. Tissue fragments (cut into 3-5 mm³) underwent primary fixation in 3% glutaraldehyde and epoxy resin embedding. Ultrastructure of endothelial of spiral artery were observed by transmission electron microscope.

### Serum Na^+^ concentration measurement

Blood collection was performed via retroorbital plexus phlebotomy under isoflurane anesthesia condition. Whole blood was allowed to clot at 4°C for 30 min, followed by centrifugation at 3,000 rpm for 10 min. Serum samples were stored at -80°C until last analysis.

### Decidual tissue sodium content measurement

Following retroorbital phlebotomy, D18 decidual tissues were mechanically dissected to remove blood clots and the smooth muscle layer. After triple washing with deionized H₂O, specimens were desiccated in a heated dry bath (37°C) until constant weight. Each dried tissue sample was digested with 1 mL concentrated HNO₃ (can add 150 µL H_2_O_2_ to help digesting). The digestate was diluted ultrapure H₂O (Milli-Q) and keep in room temperature. The last solution’s Na^+^ concentration was measured by inductively coupled plasma-optical emission spectrometry (ICP-OES, SPECTRO SPECTROBLUE FMX36). Concentration of decidual Na^+^ in the analyzed solutions were back-calculated their total content in the digested tissue samples through normalized tissue digest solution (blank) and the initial dry tissue’s weight (mmol/l).

### HUVEC culture experiments

Human umbilical vein endothelial cells (HUVECs) were maintained in endothelial growth medium (AcMec, ScienCell-1001) and incubated at 37 °C in humidified air containing 5% CO_2_. HUVECs cultured in 35-mm dishes 48 hours (cellular density about 60%-70%) and them changed into high salt medium (HSM) and urea medium (osmotic control). Briefly, HSM with additional 40 mM or 80 mM of NaCl (Sigma-Aldrich, V900058) in basic medium(49, 72–74). With the same manner, medium was supplemented with an additional urea that have an equivalent osmolality to the +40 mM NaCl medium and subsequently used on the different specified assays. Pharmacological inhibition studies were performed in the same manner, an additional 40 mM or 80 mM NaCl with or without sodium channel inhibitor of Amiloride (Acmec, Y30885) or ROCK1/2 inhibitor of Y-27632 (Selleck, S1049).

### F-actin Visualization

Cells fixed with 4% PFA were permeabilized (0.1% Triton X-100), stained with Alexa Fluor 488-phalloidin (1:200; Invitrogen A12379), and counterstained with DAPI. Z-stack images were acquired confocal laser scanning microscope (Zeiss LSM 900).

### Determination of intracellular Na^+^ content

Cell preparation: A stock solution of SBFI-AM (Santa Cruz, sc-215841) mixed with equal volume of 25%(w/v) Pluronic F-127 and diluted by adding serum-free cell culture medium into 10 μM working solution. Cells were incubated with the indicator for 3 hours at 37°C, and them adding serum-free cell culture medium for further incubated for 40 mins. Washing by DPBS three times. Fluorescence obtained by exciting the dye at 340 and 380 nm and collecting the 550 nm emission.

### Statistics

All data were presented as Mean ± SEM and each experiment incorporated a minimum of three independent samples. Statistical analyses were performed by Student’s *t*-test. Graphical representations generated in GraphPad Prism 9 and statistics analyzed by IBM SPSS Statistics 26. *P* < 0.05 was considered statistically significant, \*\**P* < 0.01, \*\*\**P* < 0.001 and \*\*\*\**P* < 0.0001.

## Data Availability

The RNA-seq generated in this study have been deposited in the National Center for Biotechnology Information Sequence Read Archive under accession number under accession number GSE304465 (token: uvqrgucctdqrxyp).

## Author contributions

Lin Huang performed the investigation and wrote the manuscript of this work. Ziying Huang provided analysis for RNA-seq. Min Li, Ning Guo, Xinran Yu and Minshan Huang provided technique help. Dunjin Chen, Haibin Wang, Shuangbo Kong and Jingsi Chen directed the research.

## Acknowledgements

This work was supported by National Natural Science Foundation of China (82288102 to H.W., 82222026 to S.K.), the National Key Research and Development Program of China (2022YFC2702500 to H.W., 2022YFC2704500 to S.K.). Natural Science Foundation of Guangdong Province (2022A1515012405; 2025A1515012340) and Guangzhou school (college) enterprise joint qualification project (2023A03J0380).

